# Dynamic growth trajectories distinguish bacteriostatic and bactericidal antibiotics at subinhibitory concentrations

**DOI:** 10.1101/2025.06.14.659040

**Authors:** E. Vaisbourd, D. S. Glass, Y. Yang, A. Mayo, A. Bren, U. Alon

## Abstract

Subinhibitory antibiotic exposures are common in clinical and environmental contexts, yet their effects on bacterial growth dynamics remain incompletely understood. We studied the temporal response of *Escherichia coli* to a panel of bactericidal (“cidal”) and bacteriostatic (“static”) antibiotics at sub-minimum inhibitory concentrations (sub-MIC). We uncover a sharp dynamical distinction between the two classes. Bacteriostatic antibiotics reduce the initial growth rate in a dose-dependent manner, similar to a nutrient starvation response. In contrast, bactericidal antibiotics do not alter initial growth rates - cells continue to grow as fast as untreated cells - until an abrupt slowdown in growth rate. The onset of slowdown occurs earlier with increasing dose, suggesting a damage accumulation mechanism leading to a lethal threshold. Cidals also show a steeper dose-response curve. We propose that bacteria respond to cidal antibiotics with a "grow-fast-then-crash" strategy that is adaptive for transient lethal threats, whereas static antibiotics trigger stress adaptation and slower growth. While clinical outcomes of statics and cidals may be similar at full inhibitory doses, these sub-MIC dynamical signatures could influence resistance evolution and treatment outcomes in biofilms or partially resistant strains. Our findings offer a dynamic framework for antibiotic classification and raise new questions about how bacteria respond to sublethal antibiotic stress.

## Introduction

Antibiotics have improved health and lifespan over the past century ^1^. They can be classified in many ways, including by their origin (natural or synthetic), spectrum of activity (narrow or broad), mechanism of action, clinical role, and resistance risk ^2,3^. Mechanistically, antibiotics target diverse bacterial functions such as translation, transcription, cell wall synthesis, DNA maintenance, and metabolism ^2,4,5^.

One particularly common classification relies on treatment outcomes - bactericidal antibiotics (“cidals”), which kill bacteria, and bacteriostatic antibiotics (“statics”), which inhibit their growth^4^. In some cases, static and cidal antibiotics target the same biological process, such as translation. Although it was once thought that cidal antibiotics are clinically superior, this notion has been challenged; treatment success depends on numerous factors beyond the cidal/static distinction ^6,7^.

Despite its widespread use, the binary distinction between bacteriostatic and bactericidal antibiotics remains ambiguous, especially at subinhibitory concentrations. The formal definition hinges on the ratio between the minimal bactericidal concentration (MBC) and the minimal inhibitory concentration (MIC), which are traditionally determined by colony-forming unit (CFU) counting and turbidity, respectively. MIC is defined as the lowest concentration preventing visible growth after an overnight culture, while MBC is the lowest concentration that reduces the viable population by at least 99.9%. An antibiotic is considered cidal if MBC/MIC < 4, and static otherwise ^8^.

In practice, the classification of an antibiotic can vary depending on the bacterial species, the experimental context, or the specific strain tested. The classic methods used to define MIC and MBC may fail to distinguish between slow death and slow growth, particularly at subinhibitory concentrations, where net population change may not reflect the underlying dynamics.

Moreover, the cidal/static classification does not reveal why some antibiotics lead to bacterial death while others inhibit growth. Thus, current definitions based on MIC and MBC ratios do not capture dynamic bacterial responses, potentially limiting clinical effectiveness and our understanding of bacterial survival strategies.

A clear phenotypic distinction between cidal and static effects, especially at subinhibitory concentrations, could help reveal bacteria’s physiological strategies to survive ^9^. Measuring growth patterns over time across a range of antibiotic concentrations can provide temporal and quantitative response features that remain invisible when classifying antibiotics based on two concentrations and a single endpoint.

In this study, we explored whether bactericidal and bacteriostatic antibiotics differ in their dynamic impacts on bacterial growth at subinhibitory concentrations. By growing *E. coli* at many subinhibitory doses across a panel of 15 antibiotics, we find that cidal and static antibiotics induce qualitatively different growth trajectories. Static antibiotics immediately reduce the growth rate in a dose-dependent manner and allow bacteria to proliferate until reaching carrying capacity. In contrast, cidal antibiotics do not affect the initial growth rate but cause an abrupt reduction of growth after a dose-dependent time. We suggest that the accumulation of antibiotic-dependent damage causes the reduction of growth rate and propose a mathematical model to explain the time dynamics and dose-dependence. Furthermore, cidal antibiotics generally have a steeper dose-response curve and higher halfway points in terms of their MIC than static antibiotics. We discuss the utility of these growth responses in maximizing fitness in short versus long-term stress conditions.

## Results

### Subinhibitory antibiotics show class-specific differences in *E. coli* growth dynamics

We sought to characterize *E. coli* growth dynamics in a wide panel of antibiotics with distinct mechanisms of action and static/cidal classification (Table 1). To this end, we measured the growth curves of exponentially growing *E. coli* MG1655 at high temporal resolution (7-10 min) in a robotic system immediately following treatment with a range of subinhibitory antibiotic concentrations ^37–40^. MIC was defined as the lowest concentration at which the growth curve remained flat or failed to reach an optical density (OD) of 0.05 after 10 hours of treatment (see Table ST1 for MIC values). We validated the static/cidal outcomes by standard definitions, measuring the MBC/MIC ratio using plate assays ^8^ (Table ST1, see Methods).

**Table 1.** Mechanistic and phenotypic characteristics of antibiotics used in this study.

| Antibiotic | Chemical nature | Target and downstream effects | Antibacterial activity in Gram-negative bacteria |
| --- | --- | --- | --- |
| Ampicillin | β-lactam <sup>5</sup> | Penicillin-binding protein. Cell wall synthesis inhibitor <sup>5,10</sup> | Bactericidal <sup>10</sup> |
| Chloramphenicol | Phenicol <sup>5</sup> | Ribosome large subunit. Inhibits translation by preventing incoming aa-tRNA binding <sup>11,12</sup> | Bacteriostatic <sup>12</sup> |
| Ciprofloxacin | Fluoroquinolones <sup>5</sup> | DNA gyrase. Causes dsDNA breaks <sup>13</sup> | Bactericidal <sup>13</sup> |
| Doxycycline | Tetracyclines <sup>5</sup> | Ribosome small subunit <sup>5</sup>. Prevents tRNA binding to the ribosome and inhibits protein synthesis <sup>14</sup>. | Bacteriostatic <sup>14</sup> |
| Erythromycin | Macrolides <sup>5</sup> | Binds the large ribosomal subunit, blocks nascent peptide exit channel <sup>15,16</sup> | Bacteriostatic <sup>17</sup> |
| Fosfomycin | Phosphonic antibiotics <sup>18</sup> | Inhibits synthesis of peptidoglycan precursors, leading to loss of cell wall integrity and lysis <sup>5,18,19</sup> | Bactericidal <sup>5</sup> |
| Gentamicin | Aminoglycoside <sup>5</sup> | 16S rRNA of the ribosome small subunit. Causes mistranslated proteins and membrane potential destabilization <sup>55</sup>. | Bactericidal <sup>5</sup> |
| Nalidixic acid | Fluoroquinolones <sup>13</sup> | DNA gyrase. Causes dsDNA breaks <sup>13</sup> | Bactericidal <sup>13</sup> |
| Nitrofurantoin | Nitrofurans <sup>20</sup> | Nitrofurantoin is reduced by flavoproteins to highly reactive electrophilic molecules that indiscriminately attack essential macromolecules (DNA, ribosomes, cell wall) <sup>20</sup> | Bactericidal at high concentrations but bacteriostatic at low concentrations <sup>20,21</sup> |
| Rifampicin | Rifamycin | rpoB (β-subunit) of the RNA polymerase. Inhibits transcription by steric blocking of elongating RNA <sup>22</sup> | Bactericidal <sup>23,24</sup>.<br>Not affecting *E. coli* <sup>25</sup><br>Static, with higher MIC for Gram-negative than Gram-positive <sup>26</sup><br>Static <sup>27</sup> |
| Spectinomycin | Aminoglycosides <sup>5</sup> | Ribosome small subunit. Inhibits the translocation of mRNA and tRNA. Does not cause mistranslation <sup>5,28</sup> | Bacteriostatic <sup>5</sup> |
| Streptomycin | Aminoglycoside <sup>5</sup> | Ribosome small subunit <sup>29</sup>. Causes mistranslation by tRNA mismatching and protein aggregates <sup>5,30,31</sup> | Bactericidal <sup>31</sup> |
| Tetracycline | Tetracyclines<sup>5</sup> | Ribosome small subunit. Tetracycline inhibits translation by preventing the binding of aminoacyl-tRNA to the A site <sup>32</sup> | Bacteriostatic <sup>12,14</sup> |
| Thiolutin | Dithiolopyrrolones <sup>33</sup> | Inhibitor of mRNA synthesis (chain elongation) <sup>34</sup> | Bacteriostatic <sup>24,33</sup> |
| Trimethoprim | Trimethoprim <sup>35,36</sup> | Dihydrofolate reductase (folA). Inhibits folate synthesis | Bacteriostatic <sup>35</sup> |

The OD traces show two distinct behaviors. The seven static antibiotics show log-linear curves with slopes that deviate from the untreated curve at very early time points in a dose-dependent manner (schematic representation in Fig. 1A; data in Fig. 1 B-H). At long times biomass reaches the carrying capacity (see SF1).

**Figure 1.**
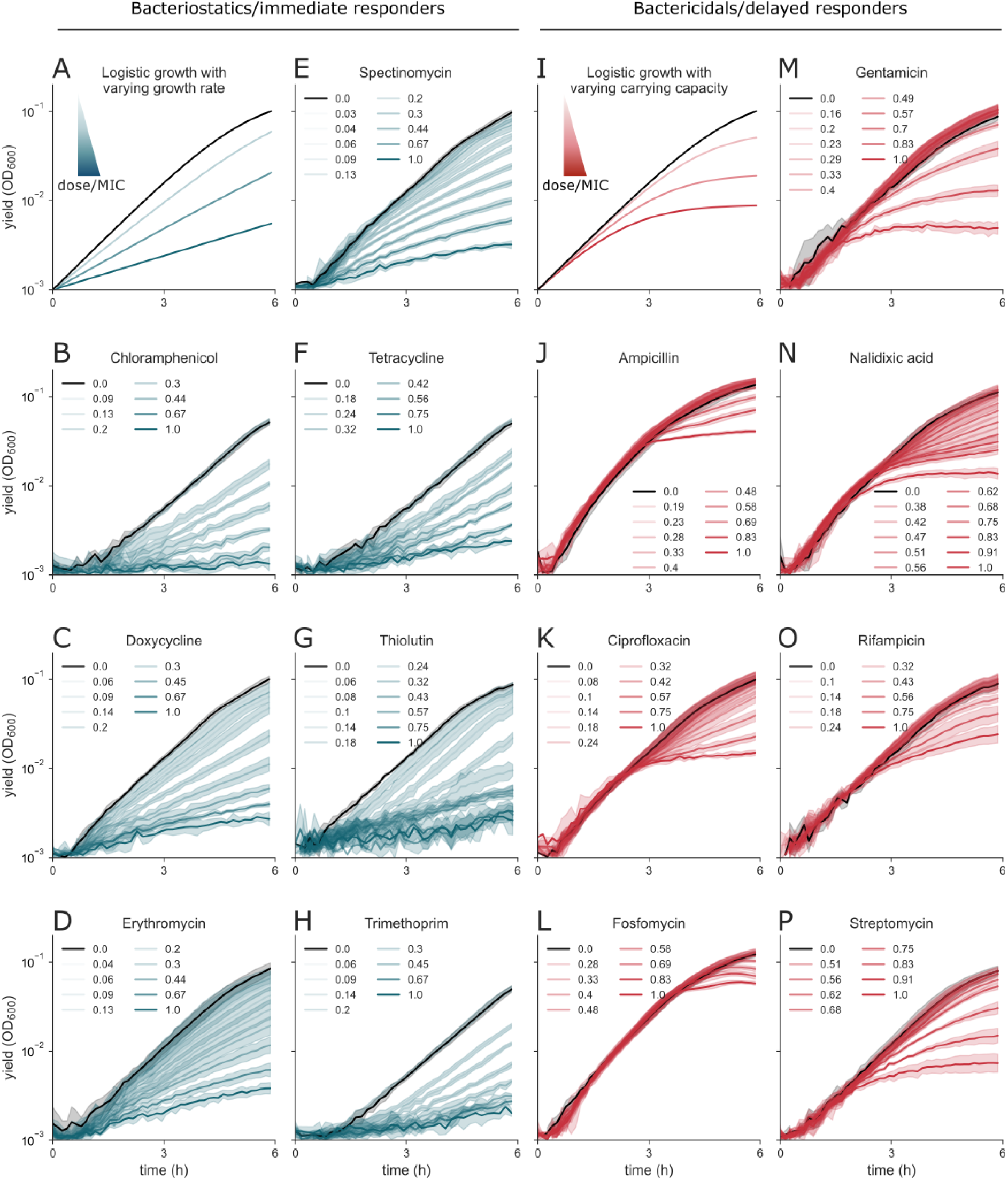
The growth traces of *E. coli* under bacteriostatic antibiotics are distinct from those under bactericidal treatment. **A, I.** Schematics showing the effects of bacteriostatic and bactericidal antibiotics on *E. coli* growth rate and yield, respectively. **B-H.** Growth curves under bacteriostatic treatment. **J-P.** Growth curves under bactericidal treatment. Treatment concentrations are normalized relative to the MIC and expressed as multiples of the MIC (Table ST1). Data are presented as mean ± SD of 5-6 technical replicates. See SF2 for two more biological replicates.

The seven cidal antibiotics show more complex growth dynamics. The growth curves are very similar to the untreated curve for the first 1.5 - 3 h, followed by a reduced growth rate that decreases in a dose-dependent manner (schematic representation in Fig. 1. I; data in Fig. 1 J-Q).

We quantified the initial growth rate (Fig. 2A,B), highlighting these two distinct phenotypes: a growth rate that drops with dose (Fig. 2C) and a growth rate that remains similar to untreated, even at doses close to the MIC (Fig. 2D).

**Figure 2.**
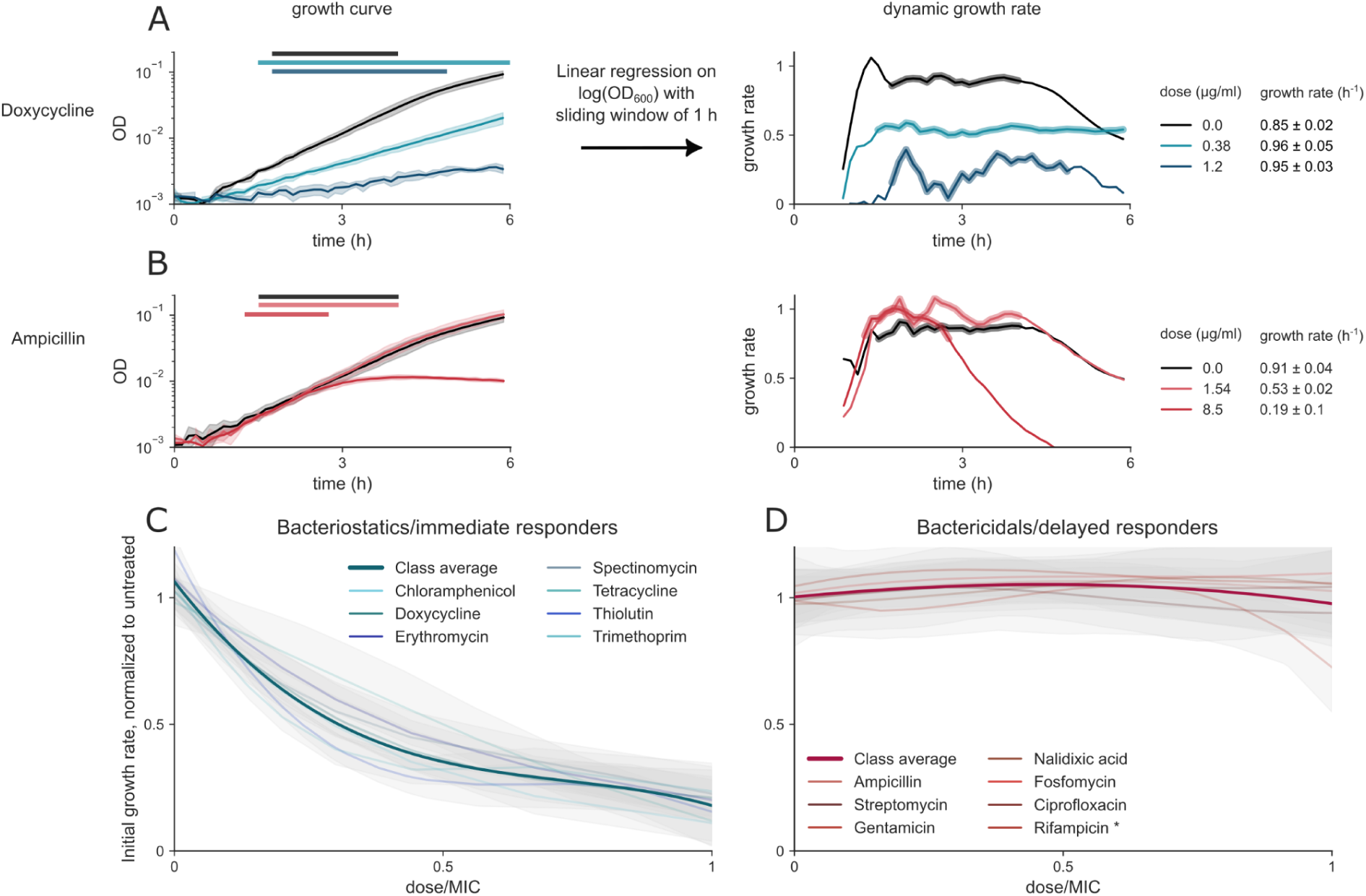
Static antibiotics reduce the initial growth rate in a dose-proportional manner, whereas cidal antibiotics do not affect the initial growth rate. **A., B.** Representative growth curves for ampicillin (A) and doxycycline (B) with horizontal bars indicating the time intervals used for growth rate estimation. In both cases, growth rates were calculated by fitting a linear regression to log-transformed OD values and averaged across the selected time interval for each treatment. **C.** Bacteriostatic antibiotics exhibit a dose-dependent inhibitory effect on growth. **D.** Bactericidal antibiotics, however, exert their effect largely independently of concentration, maintaining a similar initial growth rate to untreated controls. Rifampicin is an outlier, being a delayed responder in terms of growth rate, but bacteriostatic in the MBC/MIC assay. Antibiotic concentrations are normalized to the MIC, and growth rates are normalized to untreated cultures. Dose-response curves by drug represent spline fits across 3 biological replicates (each with 4-6 technical replicates). Class-level curves are the average of bootstrapped drug curves. See SF3 for raw data.

These two phenotypes correspond to the bacteriostatic and bactericidal classification (Table 1), with one possible exception (Rifampicin). All seven static antibiotics showed a dose-dependent drop in initial growth rate (immediate responders), and seven cidal antibiotics showed no effect on initial growth rate (Fig. 2C, D). Rifampicin was a partial exception: while its growth response was delayed in a cidal-like manner, plating assays suggested a more static-like behavior (MBC/MIC>4).

Thus, we conclude that most antibiotics can be sorted into cidal or static based on growth rate response at sub-MIC doses.

### Cidal antibiotics have steeper dose dependence and a higher halfway point than static antibiotics

We next examined the dose dependence of antibiotic effects by measuring the bacterial yield after 5 hours of treatment across a range of concentrations (Fig. 3A,B). All seven static antibiotics exhibited a gradual dose-dependent decrease in yield, consistent with a Michaelis-Menten-like behavior, characterized by Hill coefficients in the range of ∼1–2. In contrast, all seven cidal antibiotics displayed a steeper, cooperative dose-response relationship, with Hill coefficients ranging from ∼2 to 9 (Fig. 3; see also SF4).

**Figure 3.**
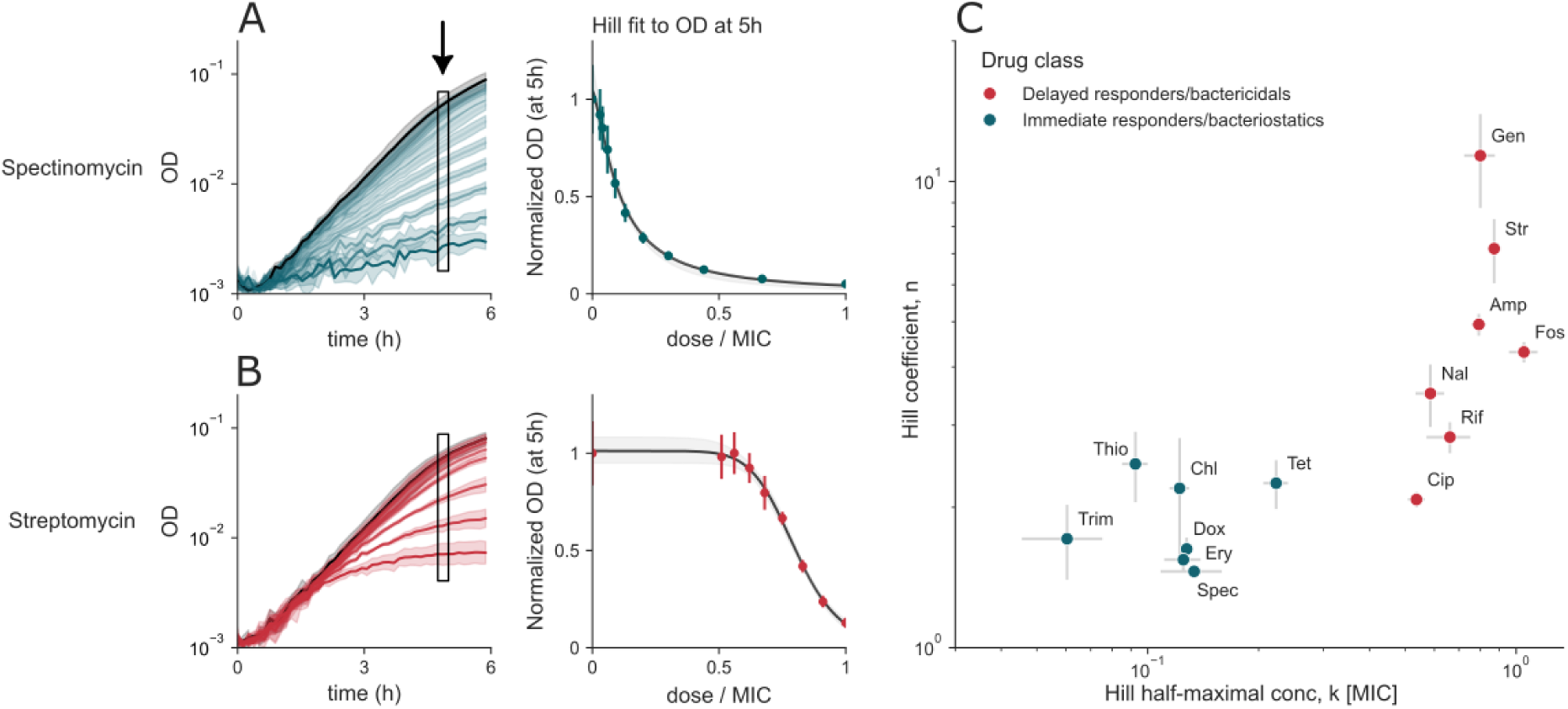
Cidal antibiotics are more cooperative and have higher halfway points than static antibiotics. **A., B.** Growth traces of spectinomycin (A) and streptomycin (B). Error bars denote the SD of technical replicates, and the grey shaded region around the Hill fit indicates 95% CI. **C.** Hill coefficients for each drug class are separable with a wide margin and cluster together. k in units of MIC (see Methods, Table ST2), error bars show SEM of three biological repeats, each with 4-6 technical replicates. Abbreviations: Trim: trimethoprim, Dox: doxycycline, Ery: erythromycin, Spec: spectinomycin, Thio: thiolutin, Chl: chloramphenicol, Tet: tetracycline, Cip: ciprofloxacin, Rif: rifampicin, Nal: nalidixic acid, Amp: ampicillin, Fos: fosfomycin, Str: streptomycin, Gen: gentamicin.

Cidal antibiotics also exhibited higher half-maximal effective concentrations (k), expressed in units of MIC, compared to static antibiotics (Fig. 3C). These trends were robust to the sampling time: analyzing the data at 4-6 hours yielded similar Hill coefficients and halfway points (see SF5).

### Growth from different initial dilutions shows duration-dependent effects in cidal antibiotics and OD-dependent effects in static antibiotics

The onset of the abrupt growth arrest in cidal antibiotics could be driven by either bacterial density or by the duration of treatment. To distinguish these two possibilities, we compared growth using an initial inoculum size spanning a 32-fold range in bacterial density (Fig. 4A). We hypothesized that if growth is density-dependent, all cultures would reach the same optical density (Fig. 4B). In contrast, if the antibacterial effect depends on treatment duration, growth curves would bend at the same time regardless of initial density (Fig. 4C).

**Figure 4.**
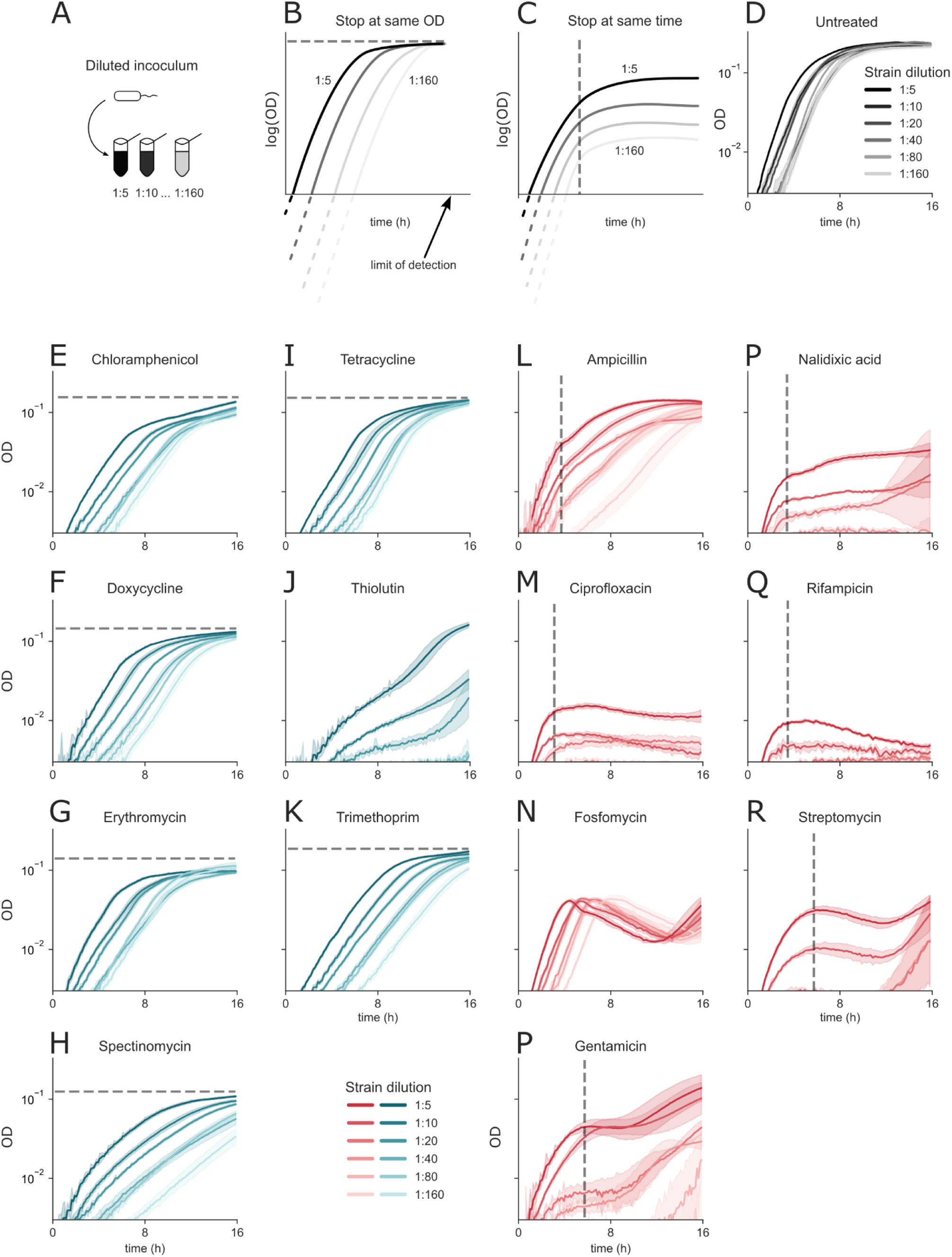
Different initial inocula show duration-dependent effects in cidal antibiotics and OD-dependent effects in static antibiotics. A. *E. coli* cultures were diluted to six inoculum concentrations (1:5, 1:10, 1:20, 1:40, 1:80, 1:160 from exponential pre-growth). **B.** Schematic representation of the expected result if growth stops at a given OD. **C.** Schematic representation of growth stops due to treatment duration. **D.** Untreated cultures reach the same OD. **E-K.** Static antibiotics stop at a given OD. **L-R.** Cidal antibiotics stop after a given treatment duration. Grey dashed lines indicate the OD- or time-dependency on the growth dynamics. Data are presented as mean ± SD of 4-6 technical replicates.

We observe that an untreated culture and cultures treated with most static antibiotics follow the same growth pattern as the density-dependent model, regardless of the initial inoculum (Fig. 4D-K). In contrast, cultures treated with cidal antibiotics growing from different inocula stop growing after approximately the same treatment duration, reaching different yields (Fig. 4L-R).

One exception to the static group is thiolutin, with growth trajectories that cannot be classified into either of the groups (Fig. 4J). Another exception is the cidal antibiotic fosfomycin, showing a time-dependent behaviour (Fig. 4N). Following the cultures over the prolonged duration (more than ∼10 hours) of treatment with cidal antibiotics reveals that some exhibit a secondary regrowth behaviour (Fig. 4N, P, R) that can be explained by mutant development or the existence of persister populations ^41^.

We conclude that the abrupt stop of bacterial growth in cidal antibiotics depends on treatment duration. We hypothesize that the time delay of the growth stop reflects the duration it takes for antibiotic-inflicted damage to become substantial enough to cause growth halt. On the other hand, growth under static treatment is limited by the growth conditions’ maximal yield (carrying capacity).

### Nitrofurantoin shows mixed growth signatures

To understand cases where the cidal/static dichotomy has difficulty in classifying antibiotics, we also studied an antibiotic that has a mixed static/cidal categorization in the literature - nitrofurantoin. It is reported to have static effects at low concentrations and cidal effects at high concentrations ^20,21^. Nitrofurantoin has a pleiotropic mechanism of action, affecting multiple metabolic and macromolecular processes in the cell ^20^ (see Table 1).

We find that nitrofurantoin shows a mixed growth signature. Growth curves in sub-MIC doses appear characteristically bacteriostatic (Fig. 5A). Yet, quantification of the initial growth rate and density at 5 h are not characteristic to either class; growth rate declines with rising dose, and the hill coefficients sit between those of the cidal and static classes (Fig. 5B-D). When cultured from varying initial inocula, we observe a mixed phenotype (Fig. 5E). When subjected to the MBC/MIC test, nitrofurantoin is lethal to cells at 4xMIC after 24 h of treatment. These observations confirm that a drug classified as mixed cidal/static also behaves intermediately in our growth-based analysis.

**Figure 5.**
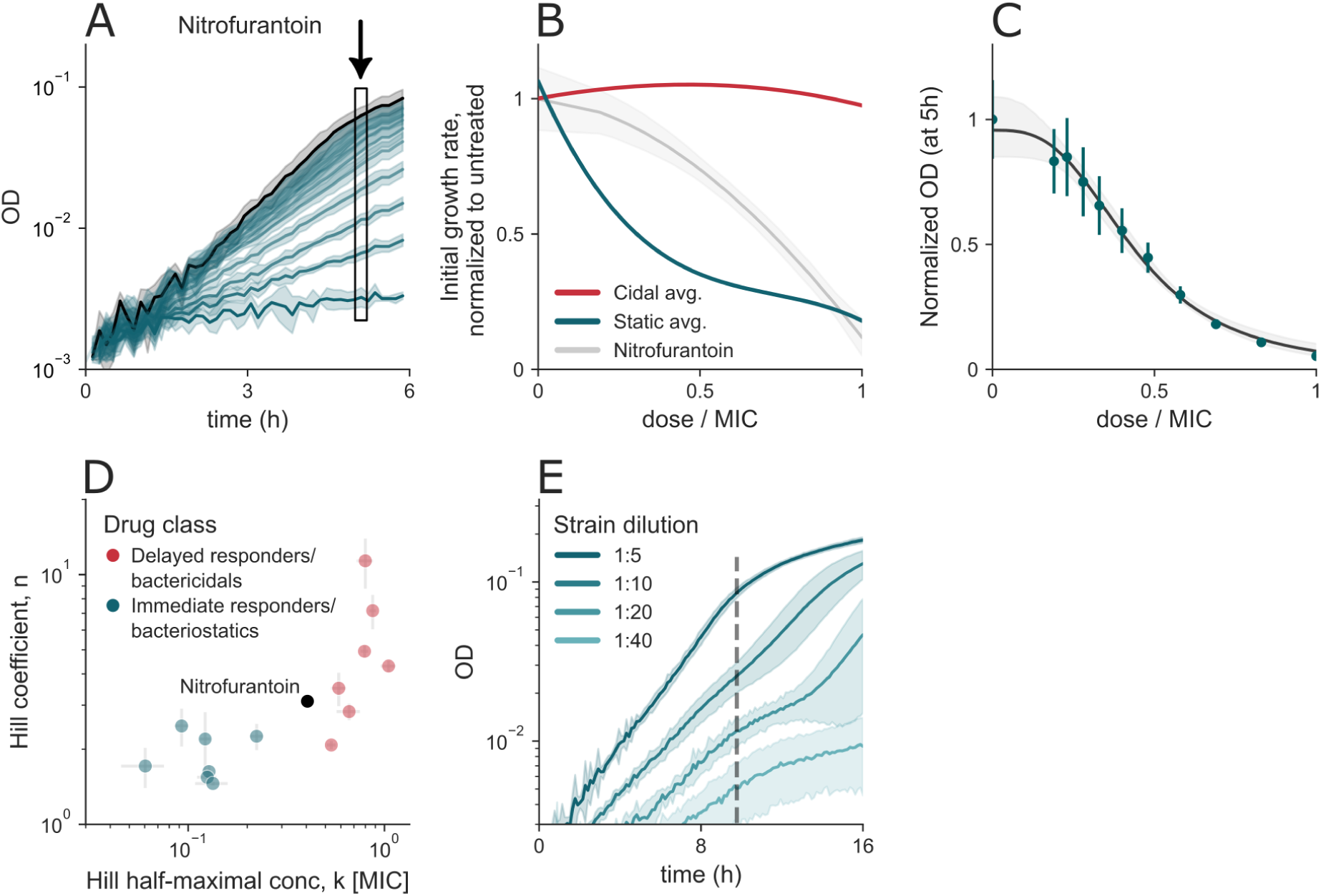
Nitrofurantoin has a mixed cidal and static signature. A. Growth curves under itrofurantoin treatment in sub-MIC concentrations. **B.** Nitrofurantoin’s growth rate decreases with treatment concentration. **C.** OD600 after 5 hours of treatment as a function of treatment concentration. Error bars denote the SD of technical replicates, and the grey shaded region around the Hill fit indicates 95% CI. **D.** Hill coefficients for nitrofurantoin fall between the cidal and static groups. **E.** *E. coli* growth under nitrofurantoin treatment exhibits both a slow growth rate and a bend, characteristics of both static and cidal antibiotics, respectively.

### Mathematical model of damage accumulation predicts the onset of slowdown in growth rate in cidal antibiotics

To establish a connection between antibiotic dose and growth dynamics, we developed a mathematical model for the abrupt slowdown in growth rate phenotype of cidal antibiotics that incorporates a latent variable, damage, as a mediating factor. Since the time to slowdown onset depends on the duration of treatment, we posit that cidal antibiotics generate intracellular damage, *y*, which stops rapid growth when it crosses a threshold 𝑦_𝑐_ ^42,43^.

We assume that the damage production rate depends on the growth rate *r*, and that damage is diluted by biomass growth. We also assume that the damage production rate is linear in antibiotic dose c. Thus, damage production rises as a function of antibiotic dose and is diluted by the growth rate *r* .

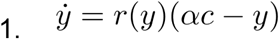

The bacterial biomass N grows with a logistic behavior, with the growth rate *r* and carrying capacity *K*.

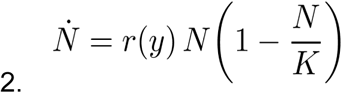

Finally, we use our observation of constant growth rate at early times to specify the dependence of growth rate on damage as constant 𝑟 when damage is below a threshold 𝑦_c_, and a Hill-like decrease with cooperativity n when damage exceeds the threshold

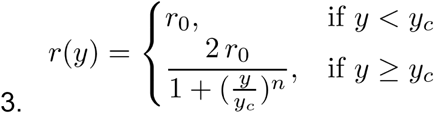

Qualitatively, the model (Eq. 1-3) captures the growth curves under cidal treatment (Fig. 6A), namely the initial exponential phase and a subsequent slowdown in growth rate.

**Figure 6.**
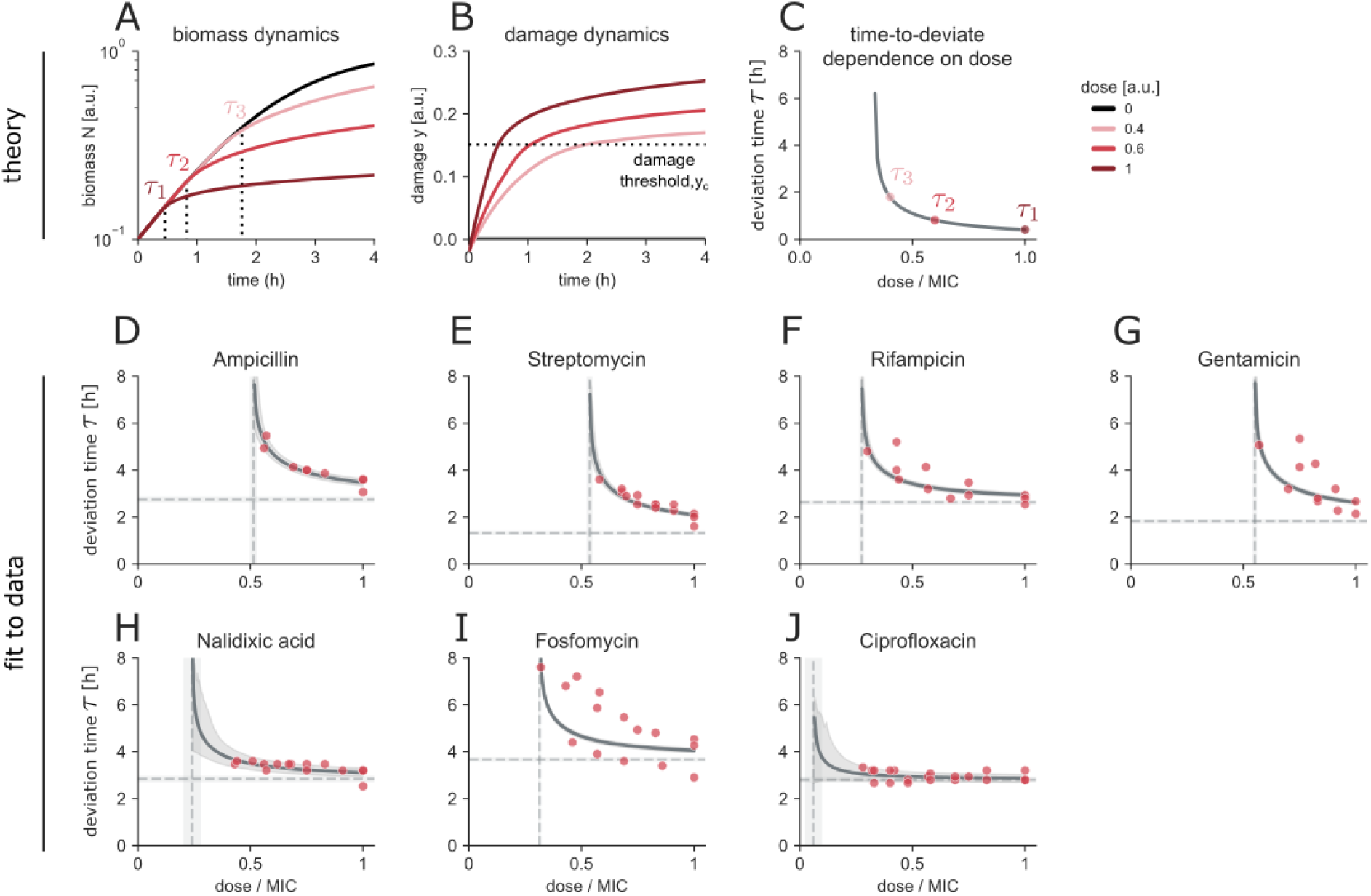
Time to deviate from the untreated curve depends on the treatment dose of cidal antibiotics and is explained by linear accumulation of damage. A. Simulated biomass accumulation dynamics of untreated (black) and treated (shades of red) bacterial cultures. Timepoints of deviation from the untreated curve are annotated τ. **B.** Simulation of damage accumulation dynamics in bacterial culture. **C.** Times of deviation as a function of the treatment dose, with annotated τ_1−3_ corresponding to 6A. **D.-J.** Fits of the model (grey) to the data (red) (N=3 biological replicates). The shaded area indicates 90% CI on the fitted parameters. Horizontal lines indicate the fitted τ_0_, the minimal duration for a deviation to occur, and the vertical lines indicate the fitted 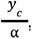, the minimal antibiotic concentration to elicit a deviation from the untreated curve.

We can determine the onset time τ when 𝑦(τ) = 𝑦_c_ by analytically solving with the initial condition 𝑦(τ_0_ ) = 0, where τ_0_ is the time needed for the antibiotic to start producing damage:

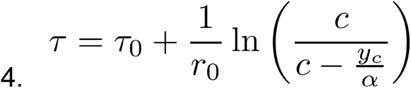

We determined the model parameters using the observed initial growth rate 𝑟_0_ and by fitting for the parameters τ_0_ and 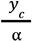 for each antibiotic. The model captured the onset of slowdown in growth rate well, with an RMS error of 0.24 h (Fig. 6D-J, ST3 for parameters and fit metrics), outperforming a baseline model (p=0.02, see Methods).

One interesting prediction is that the onset time diverges at a finite dose 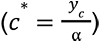, around half of the MIC for most antibiotics. Thus, there is no effect on the growth curve at low doses. This is because dilution by growth outpaces the damage production and keeps damage below the threshold. Nalidixic acid and ciprofloxacin - both DNA-gyrase inhibitors - displayed stopping times that were largely dose-independent (Fig. 6H, J), suggesting a distinct physiological response that merits further investigation.

### A tradeoff between speed and repair may shape the response

The sharp distinction between static and cidal growth responses raises a question: why do bacteria under cidal treatment continue growing rapidly and stop abruptly, rather than slowing their growth from the outset? We explore the potential evolutionary implications of this behavior through an example. Suppose that bacteria can take one of two growth strategies: grow-as-fast-as-possible or grow-slow-and-repair. In the fast strategy, cells devote a large fraction of their proteome to ribosomes, allowing rapid growth as in drug-free conditions but neglecting stress mitigation, which is detrimental in the long term. In contrast, the slow strategy allocates resources to repair or defense at the cost of a slower growth rate ^37^.

We explored these strategies under cidal and static antibiotic stresses of different durations using simulations. Brief stresses, shorter than the time to deviate τ, provide an advantage to the grow-as-fast-as-possible strategy (Fig. 7A). Longer stress periods provide an advantage to the grow-slow-and-repair strategy if at the end of the stress period the slow strategy population outnumbers the fast strategy population (Fig. 7B). One may reason that the grow-as-fast-as-possible strategy has an advantage in a situation of lethal stressors of brief duration (less than τ_0_ which is about 2-3 h in the present data).

**Figure 7.**
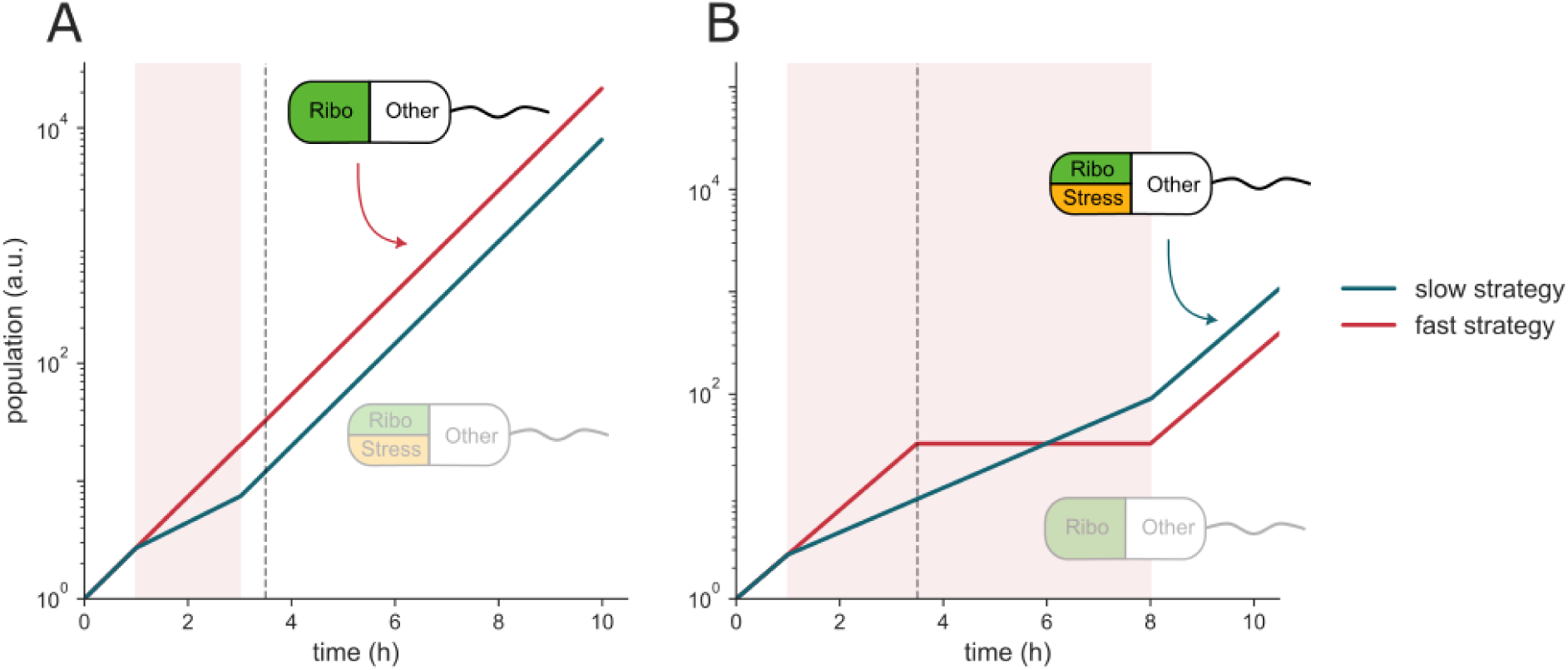
Grow-as-fast-as-possible outperforms a grow-slow-and-repair strategy when lethal stressors are brief. **A.** A short antibiotic pulse (red band, 1-4 h) ends before the deviation time τ. The ribosome-rich fast-growing population (red), therefore, accumulates more biomass than the slow, repair-biased population (teal). **B.** When the duration of stress is longer than the deviation time, damage collapses the fast population, and the slow strategy prevails. Insets schematically show how each cell partitions its resources among growth (ribosome “Ribo”, green), stress mitigation (“Stress”, orange), and other, constant functions (white); the faded cartoon marks the losing strategy in each panel. See Methods for simulation information.

## Discussion

We studied the growth dynamics of *E. coli* in response to a panel of bactericidal and bacteriostatic antibiotics at subinhibitory doses. We find a stark dynamical distinction between static and cidal antibiotics. Cidals do not affect the initial growth rate and then cause an abrupt reduction of growth, whereas statics decrease the initial growth rate in a dose-dependent way. Cidal antibiotics stop growth at a specific treatment time, which decreases with dose, and not at a fixed OD. This suggests a cell-autonomous mechanism, consistent with a mechanism in which damage accumulates until a lethal threshold is reached.

Static and cidal antibiotics show additional differences in our study. We find that OD yield drops steeply with antibiotic dose in cidal antibiotics but gradually in static ones. This is consistent with a threshold-like mechanism for damage ^8^ in cidal antibiotics. Similar threshold mechanisms were described by recent mathematical models and experiments in starving *E. coli* ^43^. Time-dependent death by cidals is consistent with ROS accumulation following nalidixic acid and ampicillin treatment that leads to death ^44^. In contrast to cidal antibiotics, static antibiotics may resemble starvation-like regulation of growth, which is titrated gradually with the restriction of cellular processes by subinhibitory antibiotic levels.

The finding of unchanged initial growth rate in cidals is consistent with previous studies on specific antibiotics^24,45,46^. Similarly, the reduced initial growth rate in statics is consistent with previous studies on specific antibiotics ^24,47^. However, to our knowledge, there have not been systematic studies of this growth signature.

A poignant comparison can be drawn between two antibiotics that target the same cellular function/mechanism - translation - where one is static (spectinomycin) and the other is cidal (streptomycin). The difference in outcomes may lie in their physiological effect. The static antibiotic reduces the translation rate and causes the cells to activate a stress - or starvation-like response that reduces the growth rate ^37,48^. In contrast, the damage caused by the cidal antibiotic causes a misfolded protein response ^49^ and might lead to upregulation of repair in a way that does not lower the growth rate. When the damage accumulates above a threshold to high levels, the cells suddenly stop growing and die ^43,47^.

A possible evolutionary optimization explanation for the unchanged initial growth in cidal antibiotics is that a grow-fast-as-possible strategy has an advantage for transient lethal challenges. A slow-and-repair strategy can be more optimal if typical challenges in the environment are prolonged (longer than a few hours).

Although clinical outcomes are often similar for bacteriostatic and bactericidal antibiotics at fully inhibitory doses, the present kinetic differences at sub-MIC doses may matter clinically for resistant strains (that pump out or neutralize the antibiotic). It may also matter when bacteria form protected niches, such as biofilms, that experience sub-MIC exposures.

Future work can test mechanisms of the cidal “crash” and static growth reduction. It would be important to test how these dynamics shape therapeutic success and immune responses. The present dynamical criterion may sharpen antibiotic classification and inspire further exploration into how bacteria survive low-dose antibiotic challenges, with the hope of improving clinical outcomes.

## Acknowledgements

We thank G. Pridham and B. Shenhar for discussions on the manuscript. We also thank R. Moran, Y. Lebel, T. Levy, L. Kreindler, and the Alon lab for comments on the manuscript.

Funding was provided by the European Research Council under the European Union’s Horizon 2020 research and innovation program (grant agreement No. 856587) and the Israel Science Foundation (grant agreement No. 1966/22).

## Author contributions

Study Design, A.B, U.A., and E.V.; Execution of Experiments, E.V.; Derivation of Theory, U.A., A.B., Y.Y., D.S.G., A.M, and E.V.; Data Analysis, E.V. and A.M; Writing Manuscript, E.V. and U.A.; Editing Manuscript, E.V., U.A., A.B., D.S.G., and Y.Y.

## Declaration of interests

The authors declare no competing interests.

## List of Supplemental Information

Figures SF1-SF5. Tables ST1-ST3.

## Materials and methods

### Strain

*E. coli* (MG1655, CGSC #6300) were grown overnight in M9 minimal medium (42 mM Na_2_HPO_4_, 22 mM KH_2_PO_4_, 8.5 mM NaCl, 18.5 mM NH_4_Cl, 2 mM MgSO_4_, 0.1 mM CaCl) with 0.05% casamino acids and 0.2% w/v glucose (M9C + glucose) at 37°C 250 rpm. For re-entry into the exponential growth phase, overnight cultures were diluted 1:300 into the same media and grown for 3-4 additional hours before treatment and initiation of measurements.

### Antibiotics

Fifteen antibiotics were used: nalidixic acid, nitrofurantoin, trimethoprim, streptomycin, spectinomycin, chloramphenicol, thiolutin, gentamicin, ampicillin, tetracycline, doxycycline, rifampicin, fosfomycin, erythromycin, and ciprofloxacin (Sigma-Aldrich). All antibiotics were dissolved in water and sterile-filtered except for trimethoprim and thiolutin (DMSO) and rifampicin (methanol). Stock solutions of antibiotics were stored at -20°C.

### Growth curve records

#### Growth under treatment with varying treatment concentrations

Antibiotics were diluted to 10 times the desired experimental concentrations manually or using a robotic liquid handler (FreedomEvo, Tecan), and 20 ul were distributed into 96-well plates by the robot. For each drug and concentration, 5-6 technical replicates were performed. Exponential phase cultures were diluted 1:10, and 180 ul were added to the plates. Then, 50 ul of mineral oil was added on top to avoid evaporation and transferred into an automated incubator, where they were grown at 37℃ and shaken (6 Hz) for 15-50 hours. To measure growth, the plates were moved by a robotic arm once every 7-10 minutes into a multi-well fluorometer (Infinite M200Pro, Tecan), and the optical density, OD at 600 nm, was recorded.

#### Growth from varying inocula

The same protocol was followed in the experiments recording growth curves from different inoculum numbers, except that after pre-growth, cultures were diluted in ratios of 1:5, 1:10, 1:20, 1:40, 1:80, and 1:160. The cultures were treated with: ampicillin (0.5 MIC), chloramphenicol (0.35 MIC), ciprofloxacin (0.9 MIC), doxycycline (0.4 MIC), erythromycin (0.35 MIC), fosfomycin (1.1 MIC), gentamicin (0.8 MIC), nalidixic acid (0.4 MIC), nitrofurantoin (1 MIC), rifampicin (1.3 MIC), spectinomycin (0.35 MIC), streptomycin (0.7 MIC), tetracycline (0.45 MIC), thiolutin (0.2 MIC) or trimethoprim (0.2 MIC). Culture conditions and OD recording were the same as for the growth curves.

### MIC determination

The minimum inhibitory concentration (MIC) for "immediate responders" was defined as the lowest treatment concentration at which OD600 remained below 0.005 after 10 hours. In contrast, the MIC for "delayed responders" was determined to be the lowest concentration that resulted in either a plateaued or declining growth curve.

### Antibiotic classification

To distinguish between bactericidal and bacteriostatic antibiotics, exponential-phase *E. coli* cultures were treated with each drug at 4×MIC for 24 hours. Before treatment, 100 ul of the untreated culture was plated, and at the end of the treatment, 100 ul of culture was serially diluted (10×) and plated on LB agar. The plates were incubated overnight at 37°C, and colony counts were compared before and after treatment. Antibiotics that reduced bacterial density by more than 3 log₁₀-fold were classified as bactericidal.

### Growth rate calculation

OD600 values were blanked by subtracting the average of the first four data points from the time series of the recorded values. Then, OD=0.001 was added to all blanked values to account for the presence of bacteria below the detection limit. Dynamic growth rates were calculated using a rolling linear ordinary least squares regression on the time series, with a window of 60 minutes. The growth rate for each drug and concentration was calculated manually by averaging the dynamic growth rate across a manually selected time window in which the dynamic growth rate was approximately constant. The error for each growth rate was estimated with error propagation. To account for day-to-day variation, treated bacterial growth rates were normalized to the untreated bacterial growth rate in each experiment.

### Growth rate dependence on dose

We estimated dose-response curves for each drug by applying spline interpolation to normalized initial growth rate measurements across concentrations normalized to MIC. To account for experimental variability, we performed bootstrap resampling (N=1000) at the level of biological replicates. In each iteration, replicates were sampled with replacement, and values were perturbed using normally distributed noise with a standard deviation equal to the replicate-specific error. The perturbed values were averaged across replicates per concentration, and a UnivariateSpline (s=1) was fit to the resulting curve.

For each drug, the mean interpolated curve was computed from the 1000 bootstrapped splines, and the standard deviation across bootstraps was used to define a shaded uncertainty band (±1 SD). At the drug class level, we aggregated all bootstrapped drug curves to compute the class-average spline, with shading again representing ±1 SD across drugs. This approach captures biological variation and measurement uncertainty in the estimated growth response.

### Hill fit to OD at 5 hours

We modeled the concentration-response relationship with a Hill function of the form

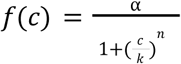

where *c* is the drug concentration, *alpha* is the maximum response, *k* is the half-maximal concentration, and n reflects the steepness. Data were aggregated by concentration and fitted using nonlinear least-squares regression. To estimate uncertainty, a nonparametric bootstrap was done, in which groups of concentration points were resampled with replacement while always including the conc=0 MIC point; the 2.5th and 97.5th percentiles of the predicted responses were computed at each concentration to yield 95% confidence intervals.

### Modeling of biomass accumulation and damage accumulation

We defined time to deviate as the time point at which the growth rate has clearly transitioned away from the untreated growth rate. These time points were manually determined by visual inspection of the plotted growth curves, with 3 biological repeats per antibiotic. Equation 4, time-to-slowdown model:

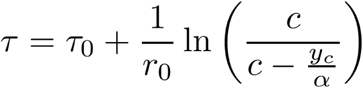

was fitted for the parameters 𝑟 and 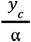 is the critical concentration (in units of MIC) below which the logarithmic term diverges, and 𝑟 = 𝑟_0_ = 1, the unchanged initial growth rate, normalized to the untreated growth rate. Parameters were estimated by non-linear least squares with bounds 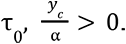. For the simulations in Fig. 6A,B (Eq.1-3), the following parameters were used: 𝑟_0_ = 1, 𝑘 = 1, α = 0. 45, 𝑦*_c_* = 0. 15, 𝑛 = 6, 𝑥_0_ = 0. 1.

### Time-to-deviate model evaluation procedure

To quantify how well the mechanism-based time-to-deviate model predicts the duration of fast growth, we compared it with a naïve mean model that simply returns the average duration observed in the training data. Evaluation was carried out independently for each antibiotic as follows. Cross-validation design: We performed leave-one-replicate-out cross-validation for each of the seven drugs: two replicates were used for training and the held-out replicate for testing, yielding three test splits per drug. We computed the mean-squared error (MSE, units = h²) for every test set between the predicted and observed duration. Using a two-sided Wilcoxon signed-rank test, we tested the null hypothesis that the median difference is zero. Our analysis showed significantly lower mean-squared error (Wilcoxon signed-rank test, n = 21, two-sided p = 0.022; significance was declared at alpha = 0.05).

### Simulation of fast and slow growth strategies

To explore how different bacterial resource allocation strategies perform under stress, we developed a simple model of population growth dynamics during a single cycle of stress followed by recovery. Two fixed strategies were considered. In the fast strategy, cells prioritize rapid growth during stress and recovery, but lack stress management. Under stress, these cells initially grow but deviate after a fixed delay τ. In the slow strategy, cells invest in stress response systems at the cost of slower growth. This strategy maintains reduced but sustained growth during stress and resumes fast growth post-stress. Simulations were implemented by integrating bacterial populations over time using:

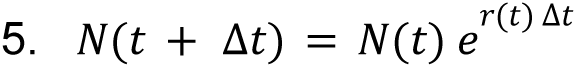

Where the growth rate r(t) depends on the current phase (stress or recovery). For both strategies, growth in stress-free conditions continues at r=1. During stress, fast strategy cells grow at r=1, whereas slow strategy cells grow at r=0.5. Each simulation began with N=1 and progressed through a drug-free phase, one stress period (2 or 7 h) with τ = 2. 5, and a recovery period. Population trajectories were calculated numerically using Euler integration with a time step Δ𝑡 = 0. 1. The population size was used as a proxy for fitness.

### Software

All calculations were performed in Python 3.10 using Numpy 2.2.4, pandas 2.2.3, and statsmodels 0.14.2. Figures were prepared in Python and edited in Inkscape.

## Notes

### Competing Interest Statement

The authors have declared no competing interest.

## References

1. Cook, M. A. & Wright, G. D. The past, present, and future of antibiotics. Sci. Transl. Med. 14, eabo7793 (2022).

2. Pancu, D. F., et al. *Antibiotics*: Conventional Therapy and Natural Compounds with Antibacterial Activity—A Pharmaco-Toxicological Screening. Antibiotics 10, 401 (2021).

3. Zanichelli, V. et al. The *WHO AWaRe (Access, Watch*, Reserve) antibiotic book and prevention of antimicrobial resistance. Bull. World Heal. Organ. 101, 290–296 (2023).

4. Baquero, F. & Levin, B. R. Proximate and ultimate causes of the bactericidal action of antibiotics. Nat Rev Microbiol 19, 123–132 (2021).

5. Kohanski, M. A., Dwyer, D. J. & Collins, J. J. How antibiotics kill bacteria: from targets to networks. Nat Rev Microbiol 8, 423–435 (2010).

6. Wald-Dickler, N., Holtom, P. & Spellberg, B. Busting the Myth of “Static vs Cidal”: A Systemic Literature Review. Clin. Infect. Dis. 66, 1470–1474 (2018).

7. Nemeth, J., Oesch, G. & Kuster, S. P. Bacteriostatic versus bactericidal antibiotics for patients with serious bacterial infections: systematic review and meta-analysis. J. Antimicrob. Chemother. 70, 382–395 (2015).

8. Pankey, G. A. & Sabath, L. D. Clinical Relevance of Bacteriostatic versus Bactericidal Mechanisms of Action in the Treatment of Gram-Positive Bacterial Infections. Clin. Infect. Dis. 38, 864–870 (2004).

9. Gil-Gil, T. et al. The evolution of heteroresistance via small colony variants in Escherichia coli following long term exposure to bacteriostatic antibiotics. Nat. Commun. 15, 7936 (2024).

10. Uehara, T., Dinh, T. & Bernhardt, T. G. LytM-Domain Factors Are Required for Daughter *Cell* Separation and Rapid Ampicillin-Induced Lysis in Escherichia coli. J Bacteriol 191, 5094–5107 (2009).

11. Jenner, L. et al. Structural basis for potent inhibitory activity of the antibiotic tigecycline during protein synthesis. Proc Natl Acad Sci U S A 110, 3812–3816 (2013).

12. Dinos, G. P. et al. Chloramphenicol Derivatives as Antibacterial and Anticancer Agents: Historic Problems and Current Solutions. Antibiotics (Basel) 5, 20 (2016).

13. Drlica, K., Malik, M., Kerns, R. J. & Zhao, X. Quinolone-Mediated Bacterial Death. Antimicrob Agents Chemother 52, 385–392 (2008).

14. Chopra, I. & Roberts, M. Tetracycline Antibiotics: Mode of Action, Applications, Molecular Biology, and Epidemiology of Bacterial Resistance. Microbiology and Molecular Biology Reviews 65, 232–260 (2001).

15. Bulkley, D., Innis, C. A., Blaha, G. & Steitz, T. A. Revisiting the structures of several antibiotics bound to the bacterial ribosome. Proceedings of the National Academy of Sciences 107, 17158–17163 (2010).

16. Menninger, J. R. & Otto, D. P. Erythromycin, carbomycin, and spiramycin inhibit protein synthesis by stimulating the dissociation of peptidyl-tRNA from ribosomes. Antimicrob. Agents Chemother. 21, 811–818 (1982).

17. Jelić, D. & Antolović, R. From Erythromycin to Azithromycin and New Potential Ribosome-Binding Antimicrobials. Antibiotics (Basel) 5, 29 (2016).

18. Raz, R. Fosfomycin: an old—new antibiotic. Clin. Microbiol. Infect. 18, 4–7 (2012).

19. Marquardt, J. L., et al. Kinetics, Stoichiometry, and Identification of the Reactive Thiolate in the Inactivation of UDP-GlcNAc Enolpyruvoyl Transferase by the Antibiotic Fosfomycin. Biochemistry 33, 10646–10651 (1994).

20. Ari, M. M. et al. Nitrofurantoin: properties and potential in treatment of urinary tract infection: a narrative review. Front. Cell. Infect. Microbiol. 13, 1148603 (2023).

21. Fransen, F., Melchers, M. J. B., Meletiadis, J. & Mouton, J. W. Pharmacodynamics and differential activity of nitrofurantoin against ESBL-positive pathogens involved in urinary tract infections. J. Antimicrob. Chemother. 71, 2883–2889 (2016).

22. Campbell, E. A. et al. Structural Mechanism for Rifampicin Inhibition of Bacterial RNA Polymerase. Cell 104, 901–912 (2001).

23. Regoes, R. R. et al. Pharmacodynamic Functions: a Multiparameter Approach to the Design of Antibiotic Treatment Regimens. Antimicrob. Agents Chemother. 48, 3670–3676 (2004).

24. Coates, J. et al. Antibiotic-induced population fluctuations and stochastic clearance of bacteria. eLife 7, (2018).

25. Hartmann, G., Honikel, K. O., Knüsel, F. & Nüesch, J. The specific inhibition of the DNA-directed RNA synthesis by rifamycin. Biochim. Biophys. Acta (BBA) - Nucleic Acids Protein Synth. 145, 843–844 (1967).

26. Wehrli, W. Rifampin: Mechanisms of Action and Resistance. Rev. Infect. Dis. 5, S407–S411 (1983).

27. Wang, Y., Fu, H., Shi, X.-J., Zhao, G.-P. & Lyu, L.-D. Genome-wide screen reveals cellular functions that counteract rifampicin lethality in Escherichia coli. Microbiol. Spectr. 12, e02895–23 (2023).

28. Mohan, S., Donohue, J. P. & Noller, H. F. Molecular mechanics of 30S subunit head rotation. Proceedings of the National Academy of Sciences 111, 13325–13330 (2014).

29. Lin, J., Zhou, D., Steitz, T. A., Polikanov, Y. S. & Gagnon, M. G. Ribosome-Targeting Antibiotics: Modes of Action, Mechanisms of Resistance, and Implications for Drug Design. Annual Review of Biochemistry 87, 451–478 (2018).

30. Busse, H.-J., Wostmann, C. & Barker, E. P. The bactericidal action of streptomycin: membrane permeabilization caused by the insertion of mistranslated proteins into the cytoplasmic membrane of Escherichia coli and subsequent caging of the antibiotic inside the cells due to degradation of these proteins. Journal of General Microbiology 138, 551–561 (1992).

31. Govers, S. K., Mortier, J., Adam, A. & Aertsen, A. Protein aggregates encode epigenetic memory of stressful encounters in individual Escherichia coli cells. PLoS Biol 16, e2003853 (2018).

32. Wilson & Daniel, N. The A–Z of bacterial translation inhibitors. Critical Reviews in Biochemistry and Molecular Biology 44, 393–433 (2009).

33. Li, B., Wever, W. J., Walsh, C. T. & Bowers, A. A. Dithiolopyrrolones: biosynthesis, synthesis, and activity of a unique class of disulfide-containing antibiotics. Nat. Prod. Rep. 31, 905–923 (2014).

34. Khachatourians, G. G. & Tipper, D. J. Inhibition of Messenger Ribonucleic Acid Synthesis in Escherichia coli by Thiolutin. Journal of Bacteriology 119, 795–804 (1974).

35. Angermayr, S. A. et al. Growth-mediated negative feedback shapes quantitative antibiotic response. Mol. Syst. Biol. 18, e10490 (2022).

36. Toth-Martinez & Bela, L. The bacteriostatic mechanisms of sulfonamidotrimethoprim combinations. Biochemical Pharmacology 26, 451–456 (1977).

37. Bren, A., Glass, D. S., Kohanim, Y. K., Mayo, A. & Alon, U. Tradeoffs in bacterial physiology determine the efficiency of antibiotic killing. Proc. Natl. Acad. Sci. 120, e2312651120 (2023).

38. Bren, A., Hart, Y., Dekel, E., Koster, D. & Alon, U. The last generation of bacterial growth in limiting nutrient. (2013) doi:10.1186/1752-0509-7-27.

39. Kohanim, Y. K. et al. A Bacterial Growth Law out of Steady State. CellReports 23, 2891–2900 (2018).

40. Zaslaver, A. et al. A comprehensive library of fluorescent transcriptional reporters for Escherichia coli. Nat Methods 3, 623–628 (2006).

41. Balaban, N. Q. et al. Definitions and guidelines for research on antibiotic persistence. Nat. Rev. Microbiol. 17, 441–448 (2019).

42. Karin, O., Agrawal, A., Porat, Z., Krizhanovsky, V. & Alon, U. Senescent cell turnover slows with age providing an explanation for the Gompertz law. Nature Communications 2019 10:1 10, 1–9 (2019).

43. Yang, Y. et al. Damage dynamics and the role of chance in the timing of E. coli cell death. Nat Commun 14, 2209 (2023).

44. Hong, Y., Zeng, J., Wang, X., Drlica, K. & Zhao, X. Post-stress bacterial cell death mediated by reactive oxygen species. Proceedings of the National Academy of Sciences 116, 10064–10071 (2019).

45. Yourassowsky, E., Linden, M. P. V. der, Lismont, M. J., Crokaert, F. & Glupczynski, Y. Correlation between growth curve and killing curve of Escherichia coli after a brief exposure to suprainhibitory concentrations of ampicillin and piperacillin. Antimicrob. Agents Chemother. 28, 756–760 (1985).

46. Comby, S., Carret, G., Pave, A., Flandrois, J. P. & Pichat, C. Bacteriostatic and bactericidal antibiotics: differences in subinhibitory concentration effects on the growth of Escherichia coli. Pathol.-Biol. 37, 335–40 (1989).

47. Comby, S., Flandrois, J. P., Carret, G. & Pichat, C. Mathematical modelling of growth of Escherichia coli at subinhibitory levels of chloramphenicol or tetracyclines. Res. Microbiol. 140, 243–254 (1989).

48. Mori, M. et al. From coarse to fine: the absolute Escherichia coli proteome under diverse growth conditions. Molecular Systems Biology 17, e9536 (2021).

49. Lindner, A. B., Madden, R., Demarez, A., Stewart, E. J. & Taddei, F. Asymmetric segregation of protein aggregates is associated with cellular aging and rejuvenation. Proc. Natl. Acad. Sci. 105, 3076–3081 (2008).

