## Supplementary Information for "Dynamic growth trajectories distinguish bacteriostatic and bactericidal antibiotics at subinhibitory concentrations"

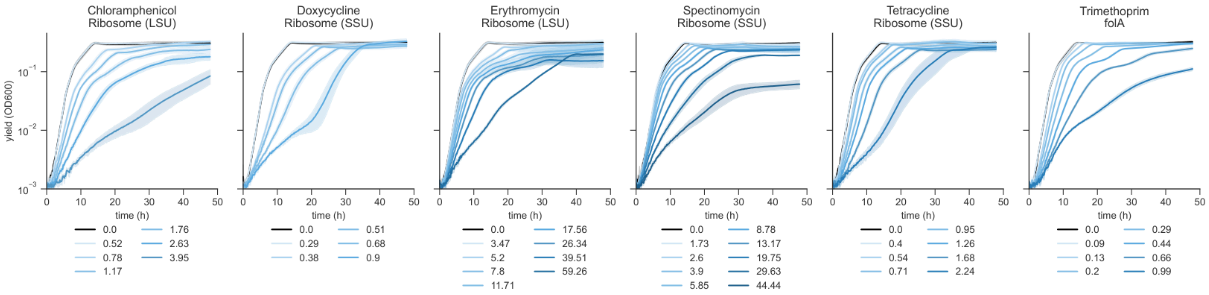


**Figure SF1. Long-term growth of *E. coli* treated with bacteriostatic antibiotics.** Exponential-phase *E. coli* were treated with sub-inhibitory doses of bacteriostatic antibiotics, and their growth curves were recorded once in 10 minutes for 48 h.

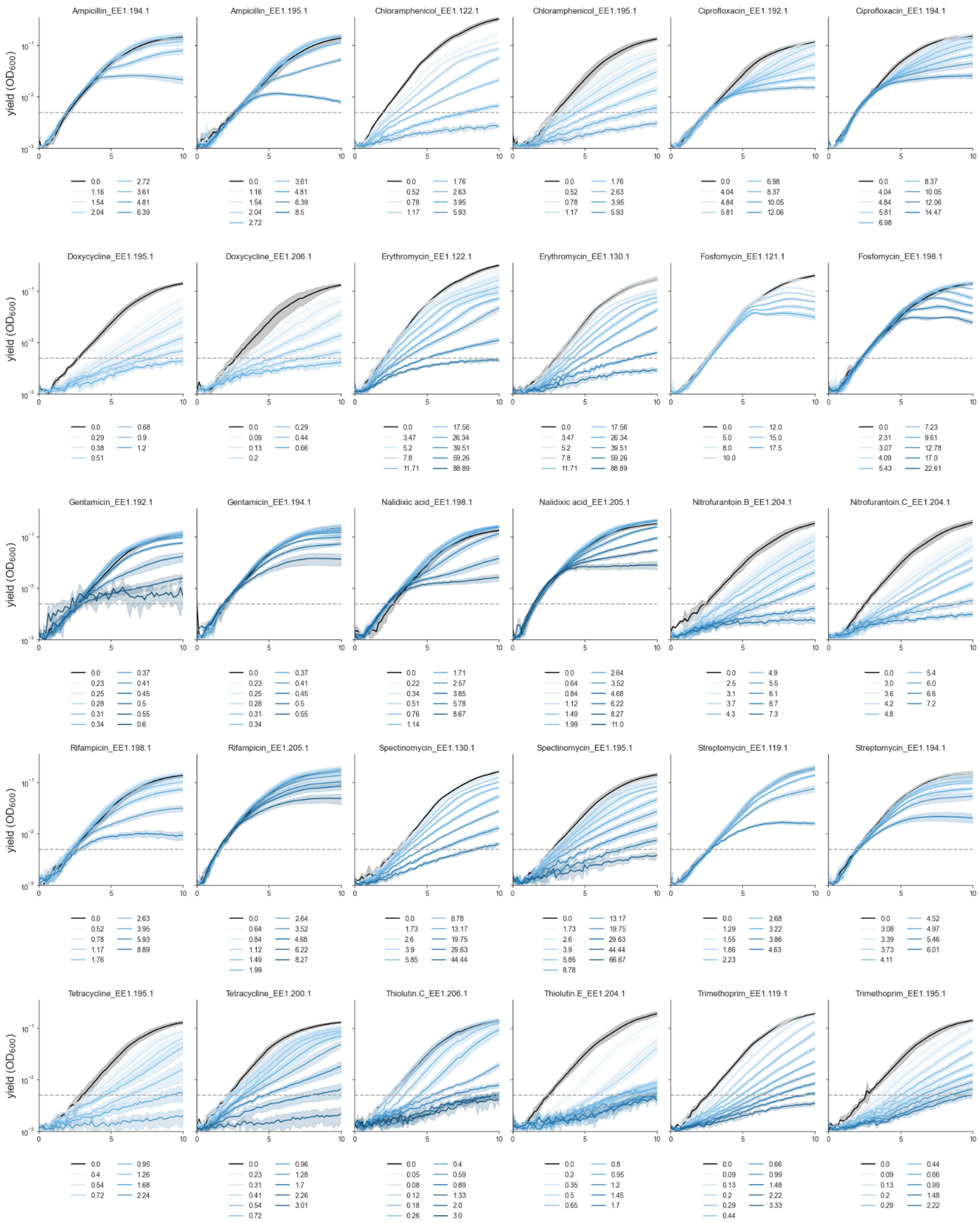

**Figure SF2. Two other biological repeats of the growth curves.**
Concentrations are indicated in µg/ml, except for ciprofloxacin, where it is ng/µl. 4-6 technical replicates per curve. Shown are treatments with concentrations smaller than or equal to the MIC in that experiment.

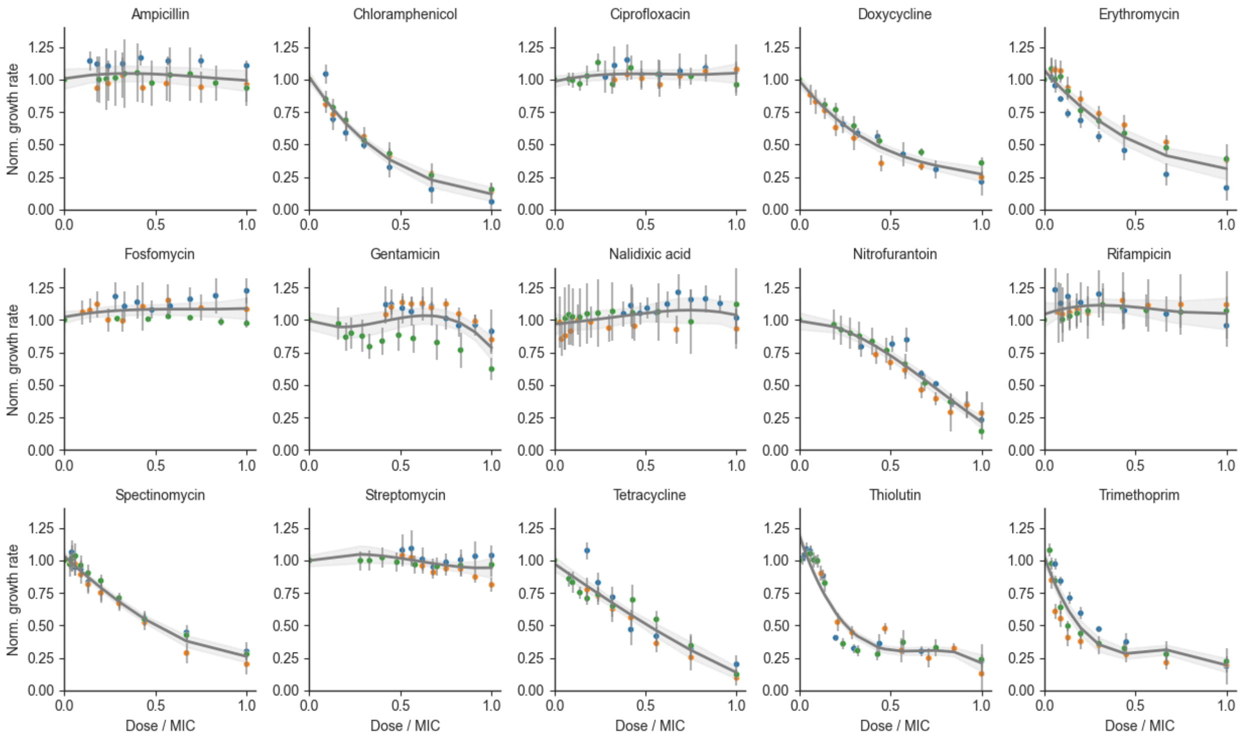


**Figure SF3. Growth rates of biological replicates.**

Growth rates are normalized to the growth rate of the untreated control, and the treatment concentration is normalized to MIC.

Each color denotes a biological replicate, and error bars denote SD across 4-6 technical replicates.

For each drug, the curve represents a spline interpolation of bootstrapped samples (N=1000), where biological replicates were resampled with replacement and measurement noise was incorporated using replicate-specific error estimates. The shaded region represents ± SD.

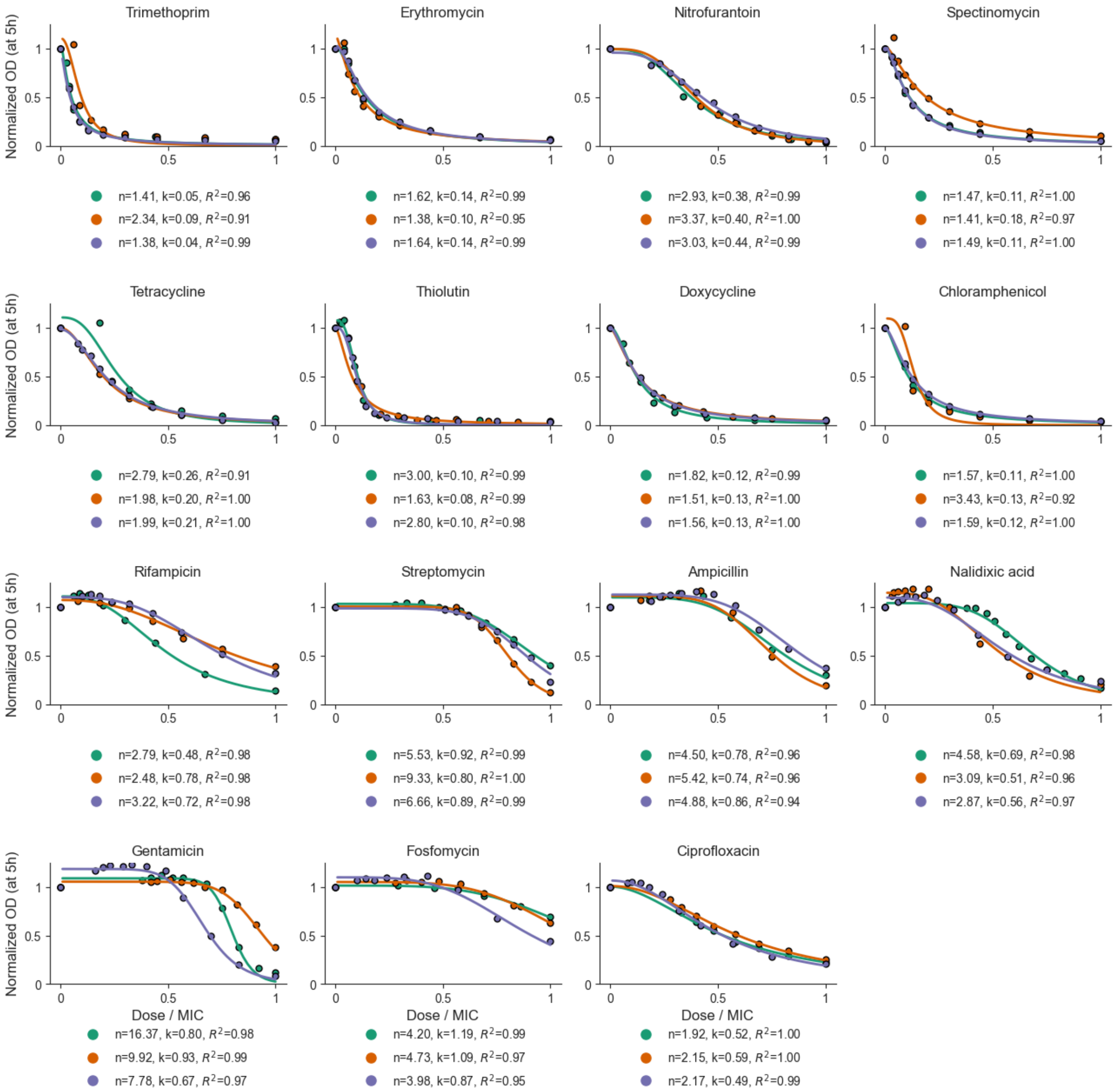


**Figure SF4. Biological repeats of the yield at 5h of treatment.** Colors indicate separate biological replicates. The fitted Hill coefficients k, n, and goodness of fit ($R^2$) are indicated for each biological replicate.

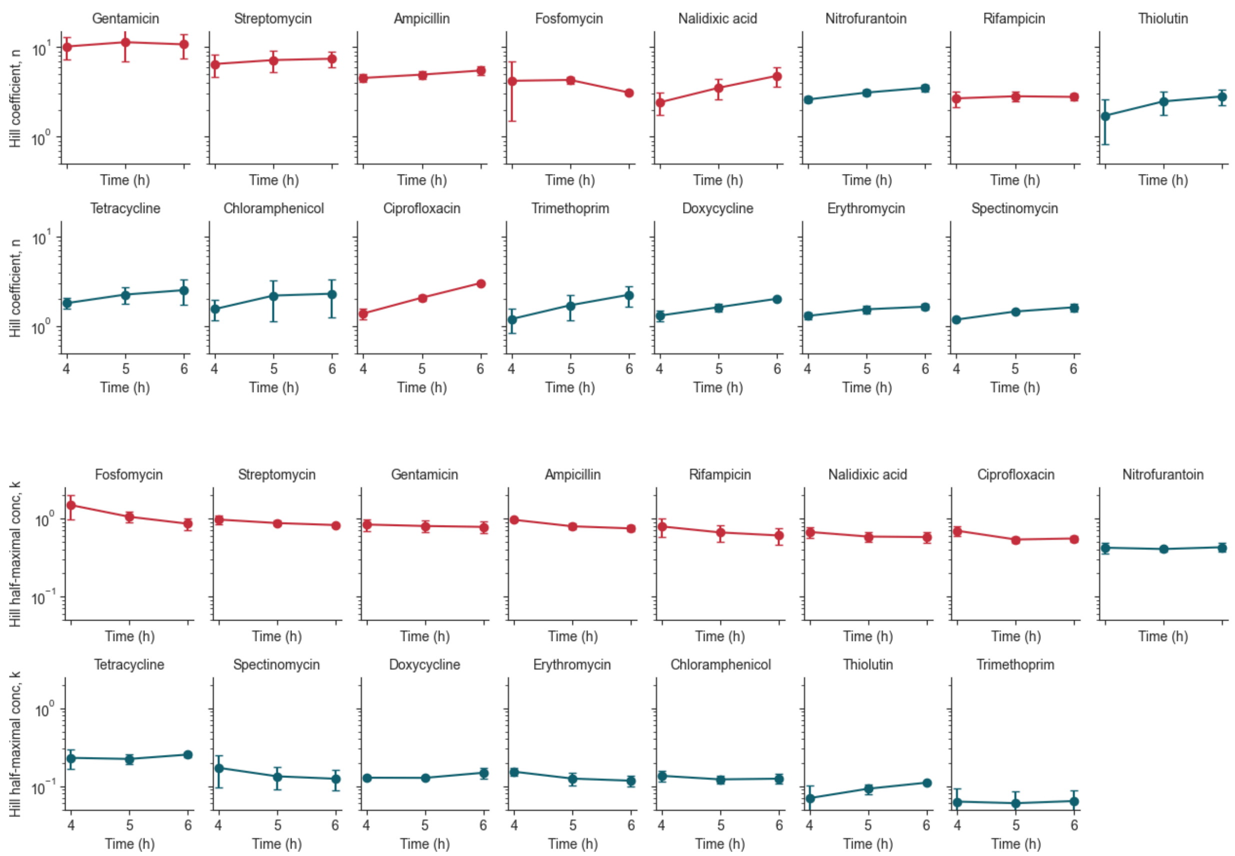


**Figure SF5. Hill coefficients n and k are insensitive to the time of sampling the growth yield across all antibiotics.** Error bars are SD of three biological replicates, each with 4-6 technical replicates. Traces are colored by drug class.

**Table ST1. Minimal inhibitory concentrations for the antibiotics used in this study.**
See Methods for the determination of MIC and the antibiotic class.

| **Antibiotic** | **MIC ± SEM** | **Growth pattern according to Fig.1** | **Antibiotic class according to MBC/MIC ratio** |
|---|---|---|---|
| Gentamicin | 0.6 ± 0.08 | delayed | cidal |
| Nalidixic acid | 8 ± 0.6 | delayed | cidal |
| Ciprofloxacin | 15 ± 3.5 | delayed | cidal |
| Ampicillin | 7.5 ± 1 | delayed | cidal |
| Fosfomycin | 18 ± 4 | delayed | cidal |
| Rifampicin | 9 ± 1 | delayed | static |
| Erythromycin | 89 ± 0 | immediate | static |
| Nitrofurantoin | 7.3 ± 0.2 | immediate | cidal |
| Streptomycin | 5.5 ± 1 | immediate | static |
| Spectinomycin | 67 ± 0 | immediate | static |
| Trimethoprim | 2.3 ± 1 | immediate | static |
| Chloramphenicol | 6 ± 0 | immediate | static |
| Tetracycline | 2.5 ± 0.5 | immediate | static |
| Doxycycline | 1.1 ± 0.4 | immediate | static |
| Erythromycin | 89 ± 0 | immediate | static |

**Table ST2. Hill coefficients for the antibiotics.**
Mean n, k, alpha ± SEM reported for 12-18 technical repeats across 3 biological repeats.

| Drug | n | k | α |
|---|---|---|---|
| Ampicillin | 4.93 ± 0.27 | 0.79 ± 0.04 | 1.12 ± 0.01 |
| Chloramphenicol | 2.20 ± 0.62 | 0.12 ± 0.01 | 1.03 ± 0.03 |
| Ciprofloxacin | 2.08 ± 0.08 | 0.54 ± 0.03 | 1.02 ± 0.02 |
| Doxycycline | 1.63 ± 0.10 | 0.13 ± 0.00 | 1.01 ± 0.00 |
| Erythromycin | 1.54 ± 0.08 | 0.12 ± 0.01 | 1.07 ± 0.04 |
| Fosfomycin | 4.30 ± 0.22 | 1.05 ± 0.09 | 1.05 ± 0.02 |
| Gentamicin | 11.36 ± 2.58 | 0.80 ± 0.08 | 1.11 ± 0.03 |
| Nalidixic acid | 3.51 ± 0.54 | 0.59 ± 0.03 | 1.10 ± 0.03 |
| Nitrofurantoin | 3.11 ± 0.13 | 0.41 ± 0.02 | 0.98 ± 0.01 |
| Rifampicin | 2.83 ± 0.22 | 0.66 ± 0.09 | 1.10 ± 0.01 |
| Spectinomycin | 1.46 ± 0.03 | 0.13 ± 0.03 | 1.03 ± 0.00 |
| Streptomycin | 7.17 ± 1.13 | 0.87 ± 0.04 | 1.02 ± 0.01 |
| Tetracycline | 2.25 ± 0.27 | 0.22 ± 0.02 | 1.03 ± 0.01 |
| Thiolutin | 2.48 ± 0.43 | 0.09 ± 0.01 | 1.03 ± 0.02 |
| Trimethoprim | 1.71 ± 0.31 | 0.06 ± 0.01 | 1.07 ± 0.01 |

**Table ST3. Fitted parameters and goodness of fit for the time-to-deviate model (Fig. 5).**
Mean fitted parameters $\tau$, $\frac{y_c}{\alpha}$ ± SD for the fits, pooled data across 3 biological repeats.

| **Drug** | $\tau\ [h]$ | $\frac{y_c}{\alpha}\ [MIC]$ | **RMSE [h]** | **MAE [h]** |
|---|---|---|---|---|
| Ampicillin | 2.7 ± 0.08 | 0.51 ± 0.01 | 0.23 | 0.19 |
| Ciprofloxacin | 2.8 ± 0.09 | 0.06 ± 0.03 | 0.21 | 0.18 |
| Fosfomycin | 3.7 ±0.06 | 0.31 ± 0.002 | 1.19 | 0.98 |
| Gentamicin | 1.82 ± 0.05 | 0.55 ± 0.01 | 0.87 | 0.60 |
| Nalidixic acid | 2.84 ± 0.1 | 0.24 ± 0.04 | 0.21 | 0.17 |
| Rifampicin | 2.63 ± 0.06 | 0.27 ± 0.01 | 0.56 | 0.38 |
| Streptomycin | 1.32 ± 0.05 | 0.54 ± 0.01 | 0.24 | 0.20 |
